# Anisotropic diffuse growth in Arabidopsis stigma papillae

**DOI:** 10.1101/2025.04.08.647848

**Authors:** Thomas C. Davis, Sharon A. Kessler

## Abstract

In angiosperms, the stigma is the first point of contact between the pollen (male) and pistil (female) during pollination. The stigma facilitates pollen capture and adhesion, compatibility responses, pollen germination, and pollen tube guidance to the transmitting tract. In *Arabidopsis thaliana*, the stigma is composed of single-celled stigma papillae that initiate from the apex of the carpels. Despite their critical function in plant reproduction, little is known about the cell and molecular mechanisms that govern stigma papillae growth and development. Using morphometric analysis of stigma papillae growth during different stages of floral development, we show that Arabidopsis stigma papillae grow via an anisotropic diffuse growth mechanism. Consistent with this conclusion, vegetative anisotropic growth mutants with defective microtubule and cellulose microfibril organization also have compromised stigma papillae growth.

## Introduction

The female gametophytes of angiosperms are buried under layers of sporophytic tissue that protect and nurture the developing seeds after fertilization. However, sexual reproduction requires that the male and female gametes be in direct contact. As a consequence, angiosperms have evolved an extraordinary tissue system, the reproductive tract, through which the male-derived pollen carries the immobile sperm through carpel tissue to the egg and central cell of the female (Johnson *et al*., 2019). The reproductive tract is a collection of highly specialized tissues from the stigma down through the style and the transmitting tract that is responsible for nurturing and guiding the pollen tube from its initiation on the stigma until the pollen tube is in close enough proximity to be attracted to an ovule for the completion of double fertilization (Crawford and Yanofsky, 2008). As the first female tissue the pollen encounters, the stigma plays a number of important roles in the sexual reproductive process including catching and adhering pollen grains, assessing pollen compatibility, initiating pollen tube growth, and guiding the pollen tube towards the transmitting tract (Edlund *et al*., 2017; Heslop-Harrison, 1978; Heslop-Harrison and Shivanna, 1977; Takayama and Isogai, 2005; Wolters-Arts *et al*., 1998).

The structure and physical properties of the stigma greatly influences its ability to complete these functions. By forming plumose, outreaching structures, many stigma are able to greatly increase their surface area to catch pollen grains from the air (Heslop-Harrison, 1981; Heslop-Harrison and Shivanna, 1977). In the model plant *Arabidopsis thaliana*, the stigma papillae are elongated, bell-shaped cells that project in all directions off of the apical surface of the style. This shape allows the stigma papillae to efficiently capture pollen grains from the nearby dehiscent anthers during self-pollination (or from pollinators in related outcrossing *Brassica* species). After receiving the signals to germinate on the stigma papillae, the pollen grain sends out a pollen tube that grows under the papillae cell wall and into the transmitting tract (Kandasamy *et al*., 1994). The basal guidance of the pollen tube on the stigma papillae is largely influenced by the mechanical properties of the papilla cell wall which in turn is dependent on the organization of cortical microtubules (CMTs) and cellulose microfibrils (CMFs) (Riglet *et al*., 2020). As stigmas age, the orientation of the CMTs and CMFs in the stigma papillae becomes more isotropic causing the pollen tube to coil around the papillae (Riglet *et al*., 2020). Despite the importance of shape and mechanical properties to stigma papillae function, little has been done to investigate the growth mechanism of stigma papillae.

Based on their elongate shapes and the expression of signaling proteins similar to tip-growth regulators in pollen tubes and root hairs, stigma papillae were proposed to grow by a tip growth mechanism (Katano and Suzuki, 2021). However, no experimental evidence was provided to support this hypothesis or to test the alternate hypothesis that stigma papillae are more like trichomes than root hairs and employ a diffuse-growth mechanism. In order to determine whether the elongated shape of stigma papillae is accomplished by tip or anisotropic diffuse growth mechanisms, we dissected the growth mechanism of stigma papillae through the use of morphometrics, subcellular markers, and genetics. Our investigation revealed that the stigma papillae are not tip-growing cells and instead grow through a diffuse growth mechanism mediated through microtubule organization. Determining the type of growth program used by stigma papillae to initiate and grow to maturity is an integral step towards understanding stigma development and plant sexual reproduction.

## Results and Discussion

### Stigma Papillae Grow in two phases

In Arabidopsis, Floral meristems sequentially initiate sepals, petals, stamens and carpels Floral staging is based on floral organ initiation and organ sizes relative to one another (Fig. S1) (Smyth *et al*., 1990). After carpel initiation at stage 6, the carpel grows as a hollow tube through stages 7 and 8. After initiation, stigma papillae then grow through stages 9-12 and reach full morphological and reproductive maturity during late stage 12 and 13. Papillae growth continues until the pollen adheres and germinates.

In order to test whether stigma papillae grow by tip growth or diffuse anisotropic growth, we first determined the growth dynamics of stigma papillae from stages 9-13 of floral development (Fig. S1). Live imaging of papillae from early stages of flower development is hampered by the need to remove the protective sepals from young flower buds. We found that exposure of developing pistils of early stage developing buds (floral stages 9-11) resulted in pistil death or failure of papillae to develop, presumably due to dehydration stress. To overcome this obstacle, we used a morphometric approach that took advantage of the developmental spectrum of floral buds in an inflorescence and used floral stage as a proxy for time. We expected that cell width would not increase over the growth phase if papillae cells are typical tip-growing cells, while changes in width would indicate diffuse growth rather than tip growth. Measuring the length and width of the papillae within each floral stage revealed that both the length and width of these cells increased dramatically over papillae development (Fig. 1). Notably, papillae width, which started at around 5.5 µm, increased to over three times its original size by stage 13 (Fig. 1B, C). This dramatic increase in cell width indicates that papillae are not undergoing classical tip growth and instead have polarized anisotropic growth (Fig. 1B), as has been suggested by the transverse arrangement of microtubules in growing papillae (Riglet *et al*., 2020). Notably, the slope of the papillae growth curve becomes steeper at around stage 12 of floral development (Fig. 1B, C). This indicates that papillae grow in 2 phases: growth in the length and width dimensions before receptivity and mainly in the length dimension after maturity if pollination is prevented.

**Figure 1:**
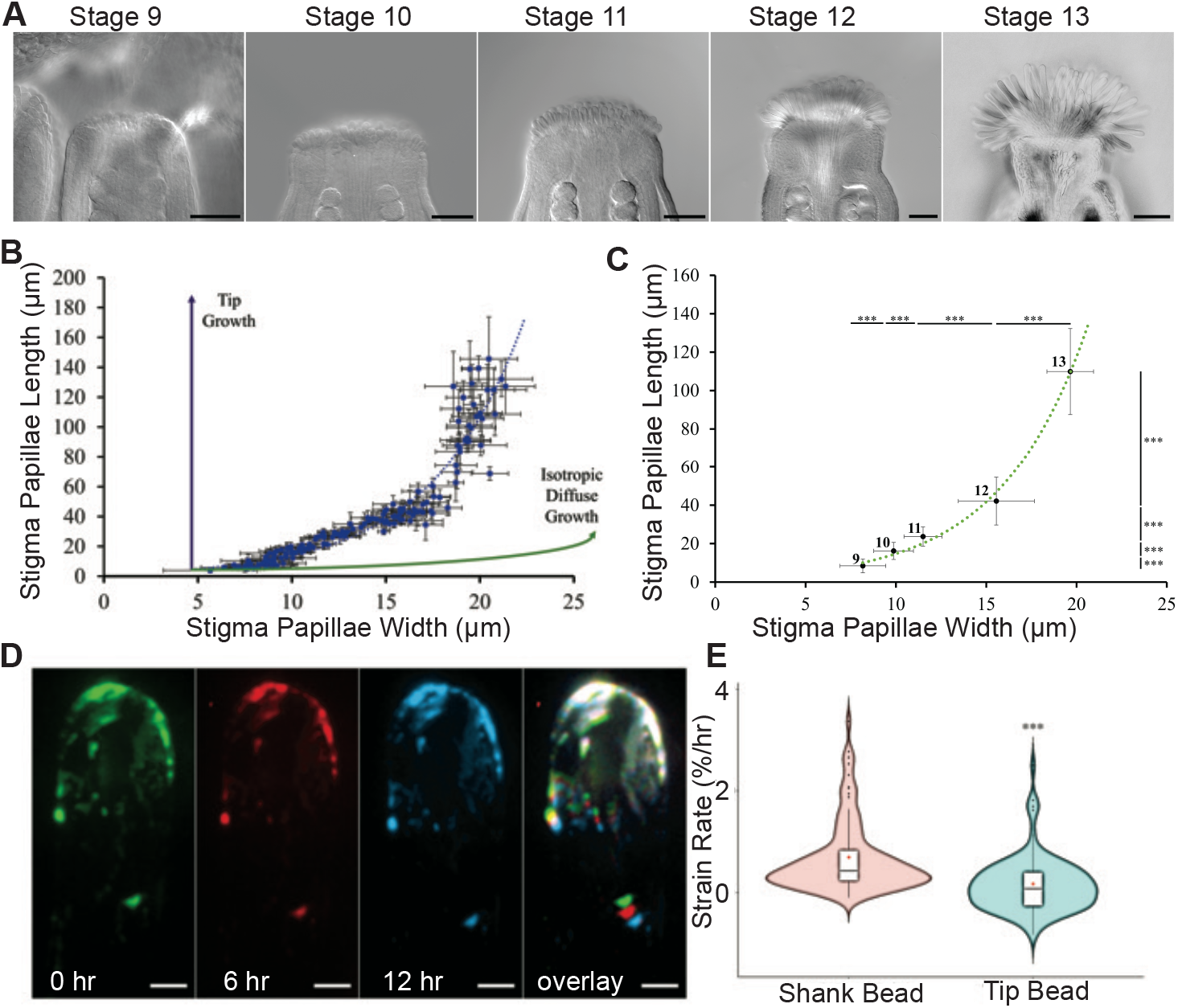
Stigma Papillae Display Anisotropic Diffuse Growth. (A) Representative images of cleared stage 9-13 stigmas are visualized using DIC microscopy. (B) Averages of stigma papillae length and widths from individual flowers are plotted against each other as blue dots with accompanying standard deviations displayed as error bars. A blue dotted line representing the exponential line of best fit is displayed. Projections of expected growth curves for tip growing and isotropic diffuse growing cells with the same starting dimensions as nascent stigma papillae are displayed as labelled purple and green lines respectively. (C) Stigma papillae length vs width for all stigma papillae from the same dataset as (B) were sorted by floral stage (labelled) and plotted. Significance between adjacent floral stages from a student’s t-test are listed in asterisk form (***: p<0.001). (D) Time lapse of growing papillae cell marked with agarose beads. (E) Strain rates of bead pairs in the shank and the tip of papillae cells. Scale bars: 50 µm (A); 5 µm (D).

To more rigorously test our diffuse growth hypothesis, we covered growing papillae from stage 12 flowers with fluorescent beads in order to measure cell wall strain over time (Elsner *et al*., 2018). We expected that tip growth would lead to cell wall strain at the tip but not shank, while diffuse growth would lead to strain throughout the cell. The fluorescent beads applied to papillae adhered to the cell wall and followed a portion of the cell wall as it grew, acting as fiducial marks (Fig. 1D). By measuring the distance between the beads every 6 hours, cell wall strain rate could be calculated as the change in distance relative to the initial distance over time (Fig. 1E). To test if papillae were diffuse growing cells, we compared the strain rate of bead pairs located 0-10 µm from the tip (tip bead pairs; n=70) against the strain rate of bead pairs located more than 10 µm from the tip (shank bead pairs; n=77) (Fig. 1E). The tip strain rate (0.17%/hr) was significantly lower than the shank strain rate (0.69%/hr) consistent with anisotropic diffuse growth where strain can be seen throughout the cell rather than just at the tip. The papillae strain rate is low compared with similarly diffuse growing cells such as Arabidopsis trichomes, which range from around 5%/hr at the base to 15%/hr closer to the tip (Yanagisawa *et al*., 2015). However, the difference in strain rate is likely a reflection of the different growth rates of fast-growing trichomes versus slow-growing stigmatic papillae.

### Stigma papillae lack the apical clear zone that is diagnostic of tip-growing cells

Tip-growing cells have a distinct subcellular organization to facilitate their growth. The majority of organelles are excluded from the tip by an actin meshwork creating what has been termed “the clear zone” (Hepler and Winship, 2015). In this zone, exocytic vesicles carrying new cell wall components are channeled radially towards the tip and endocytic vesicles carrying extra membrane are channeled centrally away from the tip in a “reverse fountain” pattern of cytoplasmic streaming (Chebli and Geitmann, 2007). We used live-imaging of stigma papillae to visualize cytoplasmic streaming. During stage 13, a mature papillae’s cytoplasm can be seen streaming around a very large central vacuole (Fig. S2A). This subcellular organization is also present during stage 12 where the majority of papillae growth occurs (Fig. S2B). There appears to be no difference between the tip subcellular organization and the shank (Fig. S2 A,B). Furthermore, by tracking subcellular structures over time, the cytoplasmic streaming pattern of stage 12 papillae suggested that, rather than a reverse fountain pattern, growing papillae stream their cytoplasm up one side of the cell and down the other (Fig. S2 C). The lack of both a clear zone and reverse fountain cytoplasmic streaming pattern are consistent with the morphometric and fiducial mark data suggesting that papillae are not tip-growing cells.

### The Diffuse Growth of Stigma Papillae Depends on the Cortical Microtubule Network

Our morphometric and fiducial mark experiments showed that stigma papillae likely grow via a diffuse growth mechanism. Diffuse growing cells rely on an orderly array of microtubules aligned transversely to the primary direction of growth (Bringmann *et al*., 2012). These microtubules act as tracks that guide cellulose synthases and ultimately the deposition of cellulose (Paredez *et al*., 2006). The cellulose microfibrils act to resist cell wall stress mostly in their direction of deposition (Green, 1980). The highly anisotropically growing papillae would be expected to have a transversely aligned cortical microtubule network. Consistent with this, transverse alignment of microtubules was observed during live imaging of stigma papillae expressing pSLR1:MAP65-citrine construct (Riglet *et al*., 2020).

Microtubule organization and cellulose biosynthesis mutants have been shown to have anisotropic diffuse growth defects in other parts of the plant. The KATANIN (KTN) and FASS1 proteins promote microtubule organization by severing microtubules that are out of alignment and promoting microtubule branch nucleation, respectively (Kirik *et al*., 2012; Nakamura and Hashimoto, 2009). Mutants in these genes grow in a dramatically more isotropic manner throughout the plant caused by their inability to organize an anisotropic diffuse growth program. The *anisotropy 1* (*any1)* mutant allele of the cellulose synthase subunit CESA1 is a non-conditional cellulose synthase mutant that produces randomly aligned cellulose microfibrils in epidermal cells and decreased crystalline cellulose leading to anisotropic growth defects throughout the plant (Fujita *et al*., 2013). Morphometric analysis of stigma papillae growth curves in these mutant backgrounds allowed us to assess the role of the cortical microtubule network and deposited cellulose on papillae growth without the limitation of starting treatment at stage 12 of floral development. In all three mutant backgrounds, papillae anisotropy was compromised (Fig. 2). For a given stigma papillae width the stigma papillae length was less than it would be for the wild type and stigma papillae were visibly wider and shorter than the wild-type papillae (Fig. 2A-H). In order to quantify and compare these differences over the course of papillae development, we transformed the growth curves for the wild-type and mutant papillae from an exponential to a linear trend using a logarithmic transformation (Fig 2I-J, Table S1, R^2^s). The dimensionless slope of the linear trend, which we termed the anisotropic growth factor (AGF), represents the degree to which the papillae length changes relative to the papillae width over the developmental time course and therefore is a good constant to measure growth anisotropy in our experimental system. All of the diffuse growth mutants had significantly lower AGFs than the Col-0 wild type (AGF=0.224±0.005), indicating that the mutant papillae grew less anisotropically (Table S1). The *any1* (AGF=0.126±0.003) and *ktn* (AGF=0.121±0.002) mutants displayed more mild phenotypes than that of *fass1* (AGF=0.095±0.002). Mature stigma papillae in the *fass1* mutant background were scarcely found as a result of cell collapse late in papillae development (Fig. 2G-H). Previous work showed that pavement cells that grow past their typical anisotropic widths are subject to increased cell wall stress (Sapala *et al*., 2018). The papillae in the *fass1* background grew far more isotropically than the cells in the *any1* and *ktn* mutants, which could have led to increased cell wall stress and subsequent collapse of the cells.

**Figure 2:**
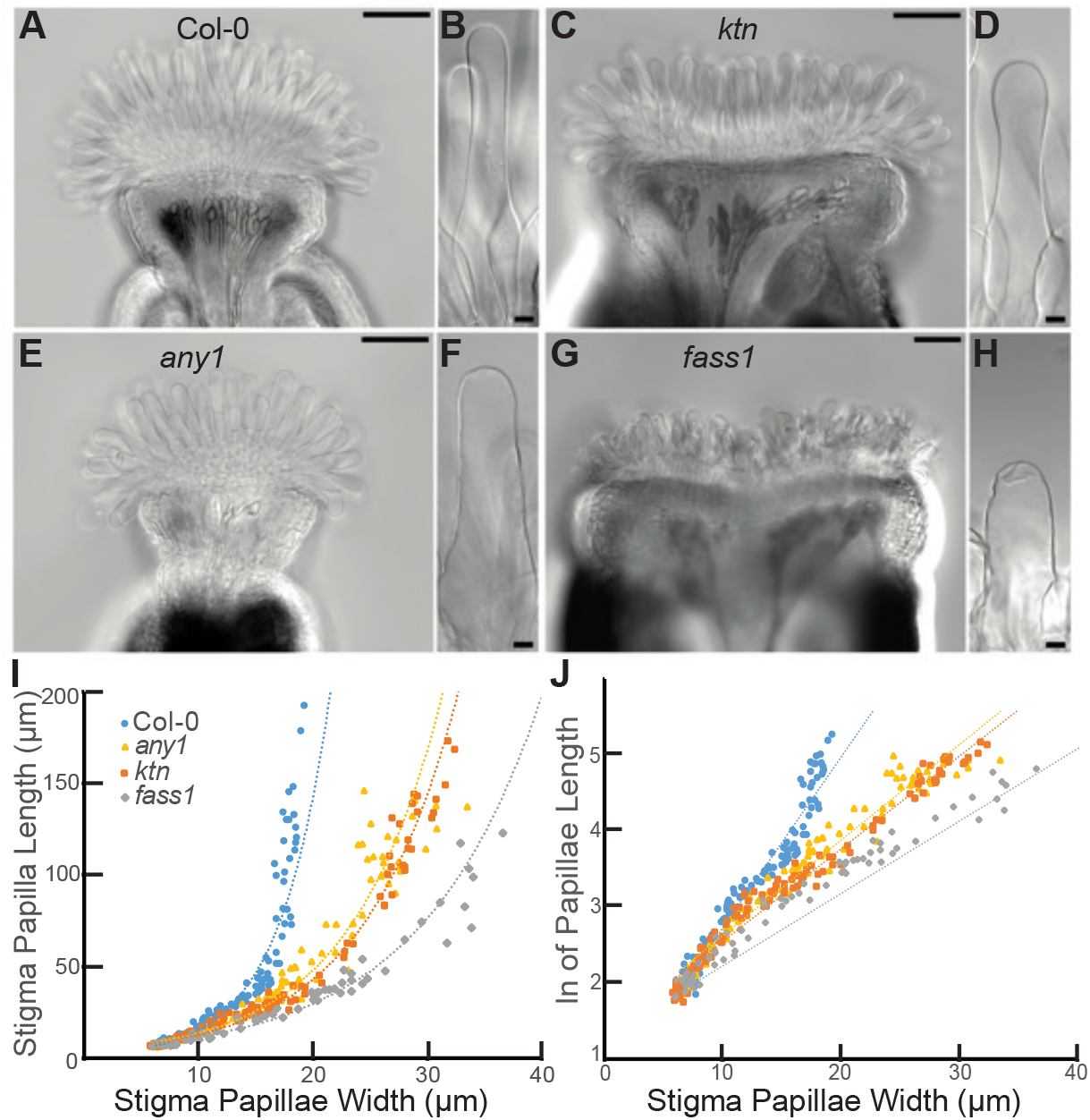
Stigma Papillae Growth is Dependent Upon Cortical Microtubule Organization and Crystalline Cellulose. (A-H) DIC images from cleared stigmas. (A) Col-0 stigma (B) Col-0 stigma papilla (C) *ktn* mutant stigma (D) *ktn* papillae cell (E) *any1* stigma (F) *any1* papillae cell (G) *fass1* stigma (H) *fass1* stigma papilla cell. (I) Papillae length vs. width growth curves for wild-type and mutant stigma papillae. Exponential lines of best fit are displayed as dotted lines. (J) natural log (ln) transformed stigma papillae lengths plotted against papillae widths. The linear line of best fit is displayed as dotted lines. Scale bars: 10 µm (A, C, E, G) and 100 µm (B, D, F, H).

Together, our results show the importance of an organized microtubule cytoskeleton and cellulose synthesis on papillae growth, which would be expected of anisotropic diffuse growing cells. These results also suggest that the papillae must achieve growth anisotropy via KATANIN, FASS1, and CESA1-dependent mechanisms in order to form the long-reaching stigma papillae that are necessary to efficiently capture pollen grains.

Our finding that papillae grow via diffuse growth rather than tip growth could be related to the function of these specialized cells during interactions with pollen tubes. Tip-growing cells typically have relatively small diameters, owing to their extremely directional growth (Campas and Mahadevan, 2009). The necessity for a tip-growing pollen tube to grow through the papillae cell wall may have placed some size minima on papillae girth forcing the papillae to grow via a growth mechanism that allowed for wider cells. Additionally, tip-growing cells tend to have highly rigid polysaccharides on their shanks to prevent radial growth that increases the width of the cell (Chebli *et al*., 2012). In contrast, the diffuse growth of papillae cell shanks likely provides variation in cell wall extensibility that is necessary to allow a pollen tube to grow into and under the papillae wall on its way to the transmitting tract. Papillae cells continue to grow after becoming receptive at floral stage 12 (see Fig 1) through senescence if they do not receive pollen. A senescence-induced programmed cell death response terminates the functional life span of the stigma papillae along with its growth (Gao *et al*., 2018).

In plants, cell size and shape are regulated by extension of the cell wall (Smith and Oppenheimer, 2005). Cell wall rigidity depends on cellulose, which is a major component of the cell wall and the extent to which it crosslinks non-cellulosic polysaccharides such as hemicellulose and pectin (Cosgrove, 2005). Cellulose is synthesized by plasma membrane-localized cellulose synthase complexes (CSCs) moving along the cortical microtubule (CMT) tracks to produce ordered cellulose microfibrils (CMFs) (Paredez *et al*., 2006). Physical forces have been shown to induce cortical microtubule reorganization in order to regulate cell shape and growth direction (Eng and Sampathkumar, 2018; Sampathkumar *et al*., 2014). KATANIN, a microtubule severing enzyme, ensures local ordering of microtubules and works downstream of auxin and Rho GTPases in this process (Lin *et al*., 2013). A role for stigmatic CMTs in pollen-papilla cell interaction was recently reported (Riglet *et al*., 2020). Riglet et al., visualized CMT organization in the stigma papillae at stages 12 to 15 of stigma development using a CMT marker MAP65.1-citrine under the control of the stigma specific SLR1 promoter. This study revealed that at stage 12 and stage 13 (at anthesis), the CMTs aligned perpendicular to the longitudinal axis of the papilla cells and were highly anisotropic. At stage 14, the CMTs were less organized and displayed higher variability in anisotropy values. At stage 15, when the stigmas extend above the anthers, CMTs displayed an isotropic orientation. The CMT organization in the papillae was correlated to pollen tube growth on the papillae. At stage 12 and 13 most of the wild-type pollen tubes grew straight into the papillae wall, while at later stages of stigma development the pollen tubes tended to coil around the papillae suggesting that the loss of CMT anisotropy impacts directional growth of pollen tube on the papillae. Consistent with this observation, *katanin1-5 (ktn1-5)* mutants have more isotropic CMT arrays in the papillae and more than 60% of the pollen tubes displayed coiling around the papillae compared to wild-type stigmas. Altered cell wall mechanical properties in *ktn1-5* papillae were linked to the resistance to pollen tube penetration (Riglet *et al*., 2020). Our study also emphasizes the importance of an organized microtubule network and cellulose synthesis machinery for anisotropic growth of stigma papillae. Most of the anisotropic growth of stigma papillae occurs by stage 13, when the stigma papillae become most receptive, allowing for compatible pollen-stigma interactions to proceed. Beyond this stage of papillae growth, the microtubule organization is more isotropic as the stigmas age and become less receptive to pollen (Riglet *et al*., 2020).

Stigma papillae are a good model system for studying the initiation, growth, and differentiation of cells that grow out of the plane of an organ. Unlike root hairs, which protrude from roots using tip growth, papillae cells grow from the apex of the pistil by anisotropic diffuse growth. Trichomes also protrude from organs using diffuse anisotropic growth mechanisms (Yanagisawa *et al*., 2015). Whether trichomes and stigma papillae have other developmental similarities is an intriguing question for future research and may lead to more insights on stigma development.

## Materials and Methods

### Plant Growth and Materials

All plants used in this study were *Arabidopsis thaliana* Columbia-0 (Col-0) ecotype. Mutant stocks used were the *katanin p60* null mutant (Elliott and Shaw, 2018) (ABRC stock number: CS816005), *fass1* (ABRC stock number: CS84613), and *any1* (Fujita *et al*., 2013; Kirik *et al*., 2012; Nakamura and Hashimoto, 2009). Seeds were sterilized in 30% sodium hypochlorite, 0.1 % Triton-X 100 for 10 minutes and a 2-3-minute incubation in 70% ethanol. Following sterilization, seeds were plated on ½ Murashige and Skoog (MS) media (Murashige and Skoog, 1962) with 0.6% phytoagar, stratified at 4 ºC for 2 days, and then moved to a light room. Seven -day-old seedlings were transplanted to soil and grown at 21-22 ºC in (16h:8h) long day conditions. Inflorescences which had already produced at least 5 stage 14 or later flowers were used in the experiments.

### Floral Staging

Floral Staging was largely based on the stages defined by (Alvarez-Buylla *et al*., 2010), but several distinct stage landmarks were used for efficient stage classification. Stage 6 was defined as the stage during which the gynoecium initiated. Stage 7 was identified by stalking of the anthers. Anther lobing was used to classify stage 8 flowers. Petal sizes of 45 to 200 µm were used as the stage 9 landmark. Stages 10 and 11 were identified by petals being longer than the short anthers but shorter than the long anthers (Alvarez-Buylla *et al*., 2010; Smyth *et al*., 1990). Style morphology was used to distinguish between stages 10 and 11. The style becomes more pronounced and style height surpasses stigma papillae length as flowers enter stage 11 and this was used as the staging landmark here. Stage 12 was defined as the stage during which the anthers were indehiscent and petals were longer than long stamens. After anthesis, flowers were marked as stage 13 flowers. Images for floral staging were taken from Col-0 inflorescences. Flowers were fixed in (9:1) ethanol: acetic acid overnight at 4 ºC. Samples were then rehydrated with a series of decreasing ethanol concentrations (85%, 70%, 50%, 30%) for 30 minutes each then placed in 50 mM phosphate buffer pH 7.0. Samples were mounted in 1:1:8 water:glycerol:chloral hydrate for clearing and were allowed to sit until optimal transparency was acquired before acquisition of images on a Nikon Ti2-E microscope equipped with 20X (N.A. 0.75) and 40X (N.A. 0.95) plan apochromat objectives and an sCMOS Monochrome camera.

### Stigma Papillae Morphometrics

With the exception of the oryzalin-treated samples (see below), all stigma papillae measurements were performed on live tissue. Carpels were first staged according to the floral staging described above then mounted in distilled water and imaged on a Nikon Ti2-E microscope with DIC. Measurements were performed using the NIS elements viewer software. Papillae that could be seen entirely in one focal plane were measured for length by using the bottom-most point and apex of the tip of the papillae. Width measurements were taken by measuring the width from both sides of the cell wall approximately 20 µm from the tip for consistency and in order to not obscure measurements by the curvature of the cell’s tip. Floral stage measurements in Fig. 1 were displayed as whole flower averages of the dimensions of 5 individually measured papillae.

### Live-Imaging Microbead Coated Papillae

Late floral stage 12 flowers were emasculated and latex polystyrene amine modified fluorescent microbeads (Sigma: L9904) in a 0.1% silwet solution were applied via a micropipettor. The bead solution was allowed to dry on the stigmas. The emasculated flowers were then excised and vortexed in a 0.4 M sucrose 3 mM MES pH 5.7 solution three times to remove any loosely attached microbeads. Samples were then transferred to ½ MS supplemented with 0.6% phytoagar contained in a gasket on a microscope slide. Samples were immersed in a 0.4 M sucrose 3 mM MES pH 5.7 solution and covered with a coverslip. Prepared samples were allowed to incubate for 1 day under standard growth conditions (see above) to allow samples to adjust to media and to minimize sample drift during imaging. After the adjustment period, samples were imaged once every 6 hours using a 60X C-Apo 1.2 NA water-immersion objective on a spinning disc confocal equipped with a CSU-10 confocal head (Yokogawa Electric) on a Zeiss Observer.Z1 system for up to 12 hours. The bead fluorophores were excited using a 561 nm laser line.

## Acknowledgements

We thank Sowmiya Devi Venkatesan for critical reading of the manuscript, the Kessler lab, Dr. Daniel Szymanski and Dr. Yun Zhou for helpful discussions, and Samy Belteton for assistance with microscopy. We thank Dr. Thierry Gaude and Isabelle Fobis-Loisy for the CMT marker line. This work was funded by startup funds from Purdue University to S.A.K.

## Supplemental Figure Legends

**Supplemental Figure 1:**
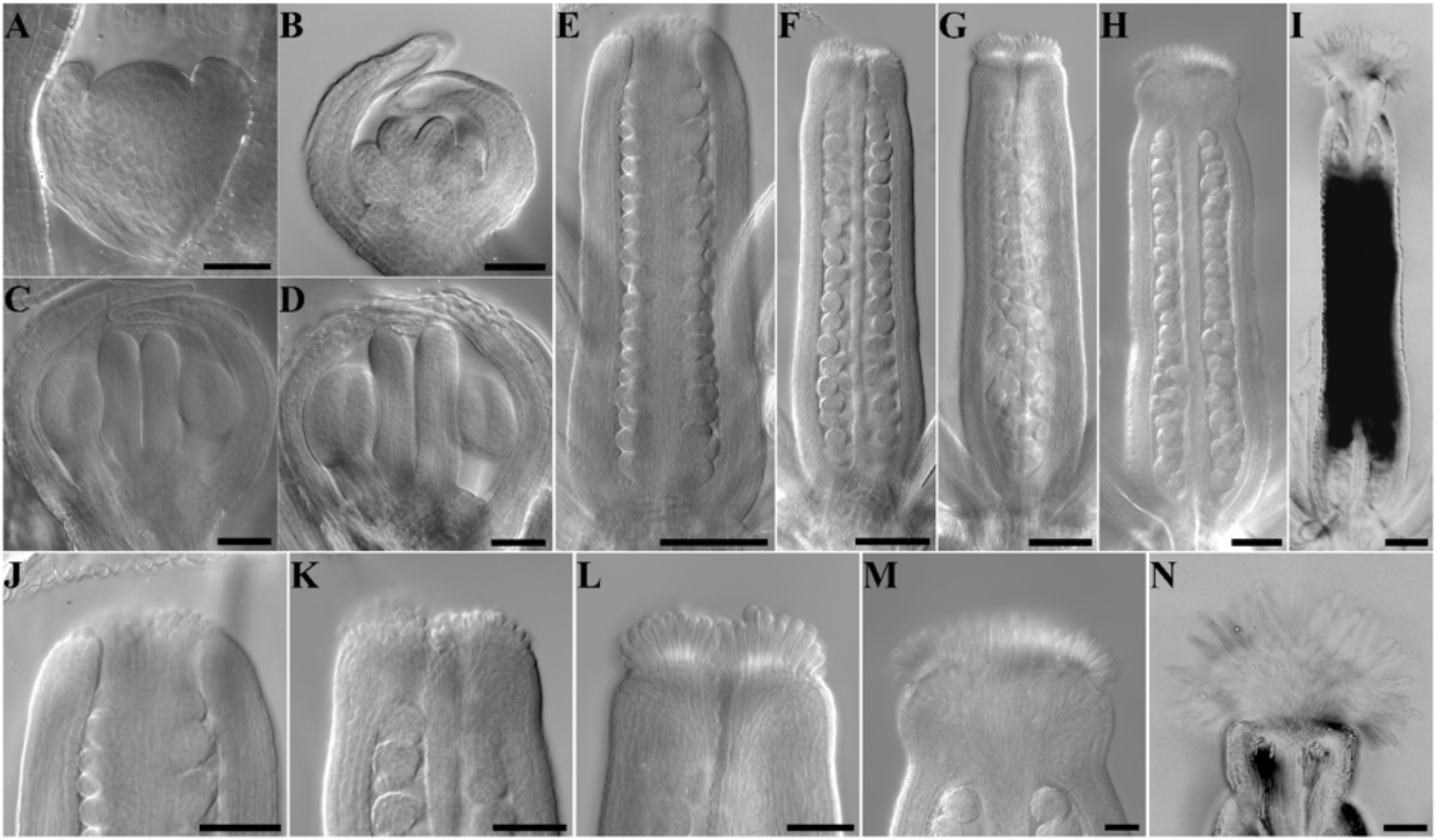
Gynoecium and Stigma Development Over Floral Stages. (A-I) DIC images of fixed and cleared gynoecia representatives from various floral stages: Stage 3 (A), Stage 6 (B), Stage 7 (C), Stage 8 (D), Stage 9 (E), Stage 10 (F), Stage 11 (G), Stage 12 (H), Stage 13 (I). (J-N) closeups of stigma from (E-I). Scale bars: 50 µm (A-D, J-K), 100 µm (E-I).

**Supplemental Figure 2:**
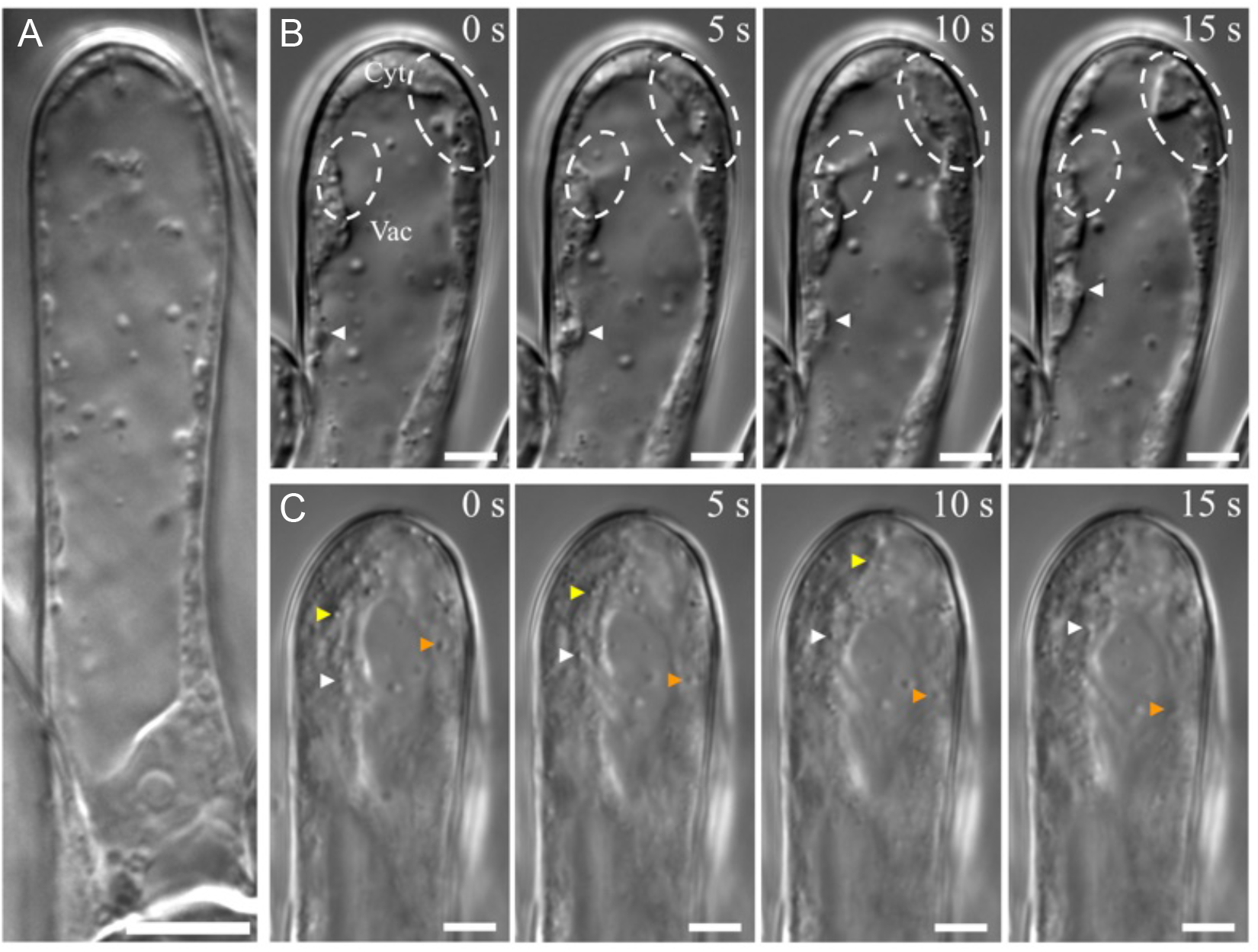
Stigma Papillae Subcellular Structure and Cytoplasmic Streaming Patterns. (A) A live stage 13 stigma papillae visualized with DIC microscopy. The vacuole (Vac) and cytoplasm (Cyt) are labelled. (B) A stage 12 stigma papillae time lapse is displayed with 5 second intervals. In the first panel, the vacuole (Vac) and cytoplasm (Cyt) are labelled. Through all panels, white ovals enclose regions where the cytoplasm can be seen moving against the vacuole and an arrowhead points to a mobile region of cytoplasm. (C) A slightly out-of-plane stage 12 stigma papillae time lapse is displayed with 5 second intervals. Different particles are tracked by different colored arrow heads over the time course to show cytoplasmic streaming. Scale bars: 10 µm (A) and 5 µm (B, C).

